# Effect of salinity on modulation by ATP, protein kinases and FXYD2 peptide of gill (Na^+^, K^+^)-ATPase activity in the swamp ghost crab *Ucides cordatus* (Brachyura, Ocypodidae)

**DOI:** 10.1101/2020.04.24.058297

**Authors:** Francisco A. Leone, Malson N. Lucena, Leonardo M. Fabri, Daniela P. Garçon, Carlos F.L. Fontes, Rogério O. Faleiros, Cintya M. Moraes, John C. McNamara

**Affiliations:** Departamento de Química, Universidade de São Paulo, Ribeirão Preto, SP; Departamento de Biologia, Faculdade de Filosofia, Ciências e Letras de Ribeirão Preto, Universidade de São Paulo, Ribeirão Preto, SP; Centro de Biologia Marinha, Universidade de São Paulo, São Sebastião, SP; Instituto de Biociências, Universidade Federal do Mato Grosso do Sul, Campo Grande, MS; Universidade Federal do Triângulo Mineiro, Iturama, MG; Instituto de Bioquímica Médica, Universidade Federal do Rio de Janeiro; Departamento de Ciências Agrárias e Biológicas, Universidade Federal do Espírito Santo, São Mateus, ES

**Keywords:** salinity acclimation, osmotic and ionic regulation, crab gill (Na^+^, K^+^)-ATPase, FXYD2 peptide, protein kinase

## Abstract

The gill (Na^+^, K^+^)-ATPase is the main enzyme that underpins osmoregulatory ability in crustaceans that occupy biotopes like mangroves, characterized by salinity variation. We evaluated osmotic and ionic regulatory ability in the semi-terrestrial mangrove crab *Ucides cordatus* after 10-days acclimation to different salinities. We also analyzed modulation by exogenous FXYD2 peptide and by endogenous protein kinases A and C, and Ca^2+^- calmodulin-dependent kinase of (Na^+^, K^+^)-ATPase activity. Hemolymph osmolality was strongly hyper-/hypo-regulated in crabs acclimated at 2 to 35 ‰S. Cl^-^ was well hyper-/hypo-regulated although Na^+^ much less so, becoming iso-natremic at high salinity. (Na^+^, K^+^)- ATPase activity was greatest in isosmotic crabs (26 ‰S), diminishing progressively from 18 and 8 ‰S (≈0.5 fold) to 2 ‰S (0.04-fold), and decreasing notably at 35 ‰S (0.07-fold). At low salinity, the (Na^+^, K^+^)-ATPase exhibited a low affinity ATP-binding site that showed Michaelis-Menten behavior. Above 18 ‰S, an additional, high affinity ATP-binding site, corresponding to 10-20% of total (Na^+^, K^+^)-ATPase activity appeared. Activity is stimulated by exogenous pig kidney FXYD2 peptide, while endogenous protein kinases A and C and Ca^2+^/calmodulin-dependent kinase all inhibit activity. This is the first demonstration of inhibitory phosphorylation of a crustacean (Na^+^, K^+^)-ATPase by Ca^2+^/calmodulin-dependent kinase. Curiously, hyper-osmoregulation in *U. cordatus* shows little dependence on gill (Na^+^, K^+^)-ATPase activity, suggesting a role for other ion transporters. These findings reveal that the salinity acclimation response in *U. cordatus* consists of a suite of osmoregulatory and enzymatic adjustments that maintain its osmotic homeostasis in a challenging, mangrove forest environment.

**Graphical abstract:** 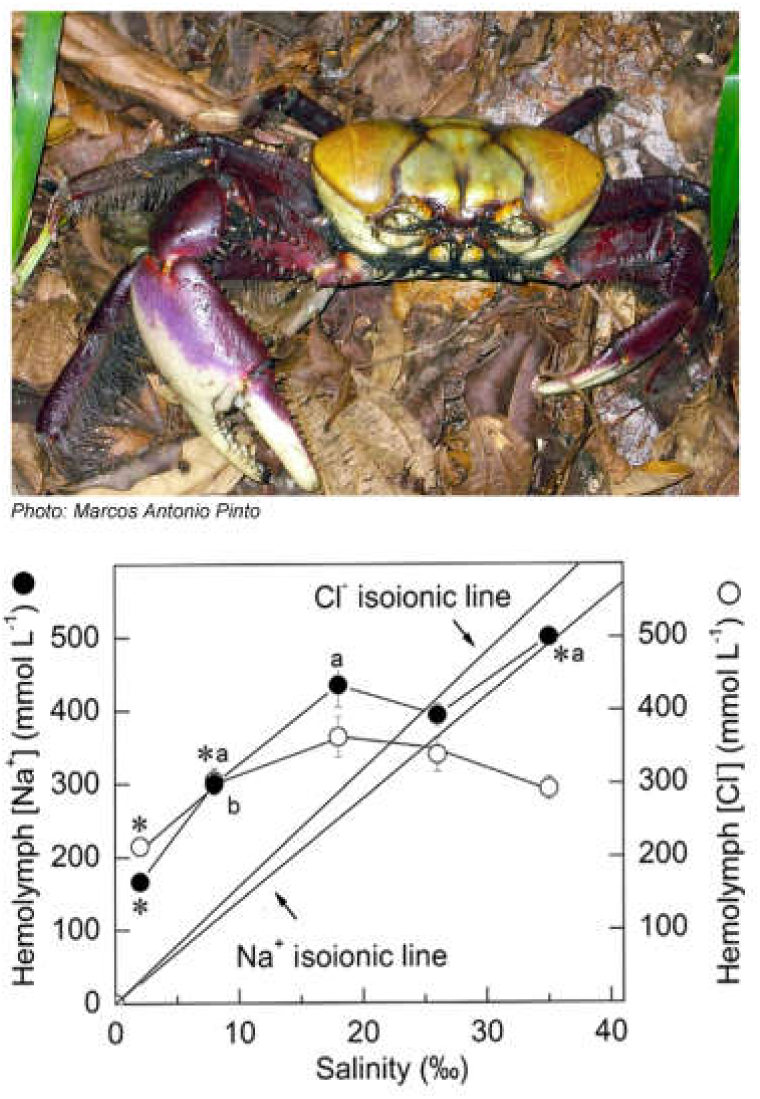

**Highlights:**

1. Gill (Na^+^, K^+^)-ATPase activity is greatest in isosmotic crabs, diminishing in lower and higher salinities.
2. A high affinity ATP-binding site (10-20% of total activity) is exposed above 18 ‰S.
3. Exogenous FXYD2 peptide stimulates activity; endogenous PKA, PKC and CaMK inhibit activity.
4. First demonstration of inhibitory phosphorylation of crustacean (Na^+^, K^+^)-ATPase by CaMK.
5. Hyper-osmoregulation shows little dependence on (Na^+^, K^+^)-ATPase activity.

## 1. INTRODUCTION

The gills, antennal glands and intestine participate in ion transport in osmoregulating crustaceans (Péqueux, 1995; Freire et al., 2008). In particular, the gills constitute a vital, multi-functional, organ effector system that contributes simultaneously to osmotic, ionic, excretory, acid-base and respiratory homeostasis (Taylor and Taylor, 1992; Péqueux, 1995; Lucu and Towle, 2003; Freire et al., 2008; Henry et al., 2012). Various enzymes including the (Na^+^,K^+^)-ATPase, V(H^+^)-ATPase and carbonic anhydrase, and ion transporters such as the Cl^−^/HCO_3_^−^ and Na^+^/H^+^ exchangers and Na^+^/K^+^/2Cl^-^ symporter, participate in the translocation of ions across crustacean gill epithelia (McNamara and Faria, 2012). Although its role in osmoregulation varies depending on the organism and its habitat, the (Na^+^, K^+^)-ATPase, particularly abundant in the cell membrane invaginations of the gill epithelial ionocytes (Towle and Kays, 1986; Taylor and Taylor, 1992; McNamara and Torres, 1999), is the main enzyme that underpins osmoregulatory ability (Lee et al., 2011).

The (Na^+^, K^+^)-ATPase is a ubiquitously expressed, integral membrane protein that couples the exchange of two extracellular K^+^ ions for three intracellular Na^+^ ions linked to the hydrolysis of a single ATP molecule (Albers, 1967; Post et al., 1972). This exchange establishes an electrochemical gradient of these ions across the plasma membrane, indispensable for many cell functions (Meier et al., 2010). The oligomeric (Na^+^, K^+^)-ATPase is a member of the P_2C_ subfamily of the P-type ATPase transporter family, and consists of an α-, a β- and a γ-subunit (Geering, 2001). The catalytic α-subunit hydrolyses ATP and transports the cations, while the β-subunit plays a crucial role in the structural and functional maturation of the α-subunit, and in modulating its transport properties (Kaplan, 2002; Morth et al., 2007). The γ-subunit is a short, single-span membrane protein belonging to the FXYD peptide family that interacts specifically with the Glu_953_, Phe_949_, Leu_957_ and Phe_960_ residues of the M9 transmembrane α-helix (Morth et al., 2007; Shinoda et al., 2009), and regulates the kinetic behavior of the (Na^+^, K^+^)-ATPase depending on cell type, tissue and physiological state (Morth et al., 2007; Geering, 2008; Shinoda et al., 2009). Its docking site on the α- subunit is highly conserved among the different ATPases, since different FXYD peptides can bind during P-type ATPase regulation (Morth et al., 2007; Geering, 2008; Shinoda et al., 2009). FXYD2 was the first FXYD protein to be linked to the (Na^+^, K^+^)-ATPase (Forbush et al., 1978). It is expressed predominantly in the mammalian kidney (Mercer et al., 1993), increases V_max_ and Na^+^ affinity without affecting ATP affinity (Cortes et al., 2006; Geering, 2006; 2008), and is a functional constituent of the *Callinectes danae* (Na^+^, K^+^)-ATPase (Silva et al., 2012). Interaction of different FXYD peptides with the (Na^+^, K^+^)-ATPase increases enzyme versatility and constitutes an important mechanism for regulating osmotic homeostasis in fish and aquatic crustaceans (Wang et al., 2008; Tipsmark et al., 2010; Yang et al., 2013, 2019a).

The (Na^+^, K^+^)-ATPase occurs in two main conformational states, E1 and E2, both phosphorylated at the D_369_ aspartate residue in which E1P shows high affinity for intracellular Na^+^ while the E2P state shows high affinity for extracellular K^+^; the binding of K^+^ accelerates dephosphorylation of the E2P form (Morth et al., 2009; Clausen et al., 2017). Analysis of the E1·AlF4^−^·ADP·3Na^+^ crystal structure from pig kidney (Na^+^, K^+^)-ATPase suggests that the M5 α-helix mediates coupling between the ion- and nucleotide-binding sites (Kanai et al., 2013). Crystallographic studies revealed either low- or high-affinity ATP binding sites present in the N-domain depending on conformational state (Morth et al., 2007; Shinoda et al., 2009; Chourasia and Sastry, 2012; Nyblom et al., 2013). The E1 conformation binds ATP with high affinity in the presence of Na^+^ (Kanai et al., 2013; Nyblon et al., 2013) while the E2 conformation binds ATP with low affinity in the presence of K^+^ (Morth et al., 2007; Shinoda et al., 2009). Na^+^-like substances such as Tris^+^ induce exposure of the high affinity ATP-binding site (Middleton et al., 2015; Jiang et al., 2017). However, despite nucleotide binding with an efficiency similar to Na^+^, the enzyme does not assume the Na^+^-like E1 form (Middleton et al., 2015).

(Na^+^, K^+^)-ATPase activity can be regulated by phosphorylation (Beguin et al., 1994; Cheng et al., 1999; Pearce et al., 2010), and both cAMP-dependent protein kinase A (PKA) and Ca^2+^-dependent protein kinase C (PKC) can phosphorylate the α-subunit (Beguin et al., 1996; Pearce et al., 2010; Poulsen et al., 2010) leading to activity inhibition. The N-terminal domain of the (Na^+^, K^+^)-ATPase is phosphorylated by PKA (Beguin et al., 1994; 1996) but the only well-characterized phosphorylation target known in the enzyme structure is the Ser_943_ residue in NKAa1, present in a short helical segment between transmembrane α-helices M8 and M9, which is a putative PKA phosphorylation site (Poulsen et al., 2010). Phosphorylation of the α1-subunit of the (Na^+^, K^+^)-ATPase by the main alpha, beta and gamma PKC isoforms leads to activity inhibition (Kazaniets et al., 2001) as does phosphorylation of rat kidney α1, α2 and α3 subunits (Blanco et al., 1998). Phosphorylation of Ser_23_ in rat (Na^+^, K^+^)-ATPase α1-transfected renal COS cells by PKC leads to intracellular Na^+^ accumulation and inhibition of both ATP hydrolysis and Rb^+^ transport (Belusa et al., 1997). However, in cells transfected with Ser23 to Ala23 α1-mutants, the (Na^+^, K^+^)-ATPase cannot be phosphorylated by PKC (Poulsen et al., 2010). Phosphorylation of rat (Na^+^, K^+^)-ATPase α-subunit by an endogenous Ca^2+^/calmodulin-dependent protein kinase (CaMK) inhibits catalytic activity significantly (Netticadan et al., 1997) and constitutes part of a mechanism mediating Ca^2+^ effects on the enzyme (Yingst et al., 1992; Lu et al., 2016).

Euryhaline crabs exhibit adjustments in gill (Na^+^, K^+^)-ATPase activity and α-subunit mRNA expression in response to salinity change (Lovett et al., 2006a; Serrano et al., 2007; Masui et al., 2009; Garçon et al., 2009; Faleiros et al., 2018). The effects of reduced salinity on gill (Na^+^, K^+^)-ATPase activities have been investigated in the blue crabs *Callinectes ornatus* (Garçon et al., 2009; Leone et al., 2015), *C. danae* (Masui et al., 2009) and *C. sapidus* (Lovett et al., 2006b; Serrano et al., 2007), the hermit crab *C. symmetricus* (Faleiros et al., 2018; and Lucena et al., 2012; Antunes et al., 2017 as *C. vittatus*) and the estuarine crab *Neohelice granulata* (Castilho et al., 2001; Genovese et al., 2004; Luquet et al., 2002; 2005). However, it is not clear whether the consequent increases in (Na^+^, K^+^)-ATPase activity result from enzyme activation, synthesis and recruitment of new protein to the cell membrane (Henry et al., 2002) or from adjustment of transport activity through regulatory phosphorylation (Silva et al., 2012).

*Ucides cordatus* (Linnaeus 1763) is a mangrove crab known as the swamp ghost crab or ‘caranguejo-uçá’ in Brazil and is one of two species of *Ucides* belonging to the family *Ucididae* (Melo, 1996). It plays an ecologically relevant role in nutrient recycling and substrate bioturbation (Nordhaus and Wolff, 2007; Nordhaus et al., 2009). The crab inhabits mangrove forests on western Atlantic Ocean shores and is distributed from Florida to southern Uruguay (Coelho and Ramos, 1972). This semi-terrestrial brachyuran exhibits a modest degree of terrestriality, absorbing water from moist substrates to compensate for loss due to desiccation and urinary excretion (Hartnoll, 1988). *Ucides cordatus* confronts substantial fluctuations in salinity, from 8 to 33 ‰S, owing to tides, frequent rain and high temperatures (Santos and Salomão, 1985a) and is a strong hyper-/hypo-osmoregulator (Martinez et al., 1999), exhibiting a hemolymph osmolality of from 700 to 800 mOsm kg^-1^ H_2_O. Salt is taken up from the external medium below 26 ‰S but is secreted in more concentrated media (Santos and Salomão, 1985a, 1985b). Hemolymph [Na^+^] ranges from 300 to 390 mmol L^-1^ in salinities above 34 ‰S (Santos and Salomão, 1985b). Acclimation of submerged *U. cordatus* to dilute seawater increases (Na^+^, K^+^)-ATPase activity by ≈1.5-fold in the posterior gills and by ≈2-fold in the antennal glands (Harris and Santos, 1993b). However, while osmoregulatory ability seems well characterized, little is known of the biochemical processes underlying ion transport in *U. cordatus* gills.

Here, we evaluate the osmotic and ionic regulatory abilities of *U. cordatus* after 10 days acclimation to hypo-, iso- or hyper-osmotic salinities, and we analyze the kinetic behavior of the posterior gill (Na^+^, K^+^)-ATPase. We also evaluate the regulation of gill enzyme activity *in vitro* by the endogenous protein kinases PKA, PKC and CaMK, and by exogenous FXYD2 peptide, aiming to further elucidate osmoregulatory mechanisms in semi-terrestrial crustaceans.

## 2. MATERIALS AND METHODS

### 2.1. Material

All solutions were prepared using Millipore MilliQ ultrapure, apyrogenic water with a resistivity of 18.2 MΩ cm. Tris (hydroxymethyl) amino methane, ATP di-Tris salt, NADH, pyruvate kinase (PK), phosphoenol pyruvate (PEP), imidazole, N-(2-hydroxyethyl) piperazine-N’-ethanesulfonic acid (HEPES), lactate dehydrogenase (LDH), sucrose, ouabain, KN62, H89, Phorbol-12-myristate 13-acetate (PMA), phosphatidyl serine (PS), bovine serum albumin, dibutyryl cAMP (db-cAMP), dithiothreitol (DTT), ethylene glycol tetraacetic acid (EGTA), chelerythrine, alamethicin, theophylline, calmodulin (CaM), thapsigargin, aurovertin, ethacrynic acid, ethylene diamine tetraacetic acid (EDTA), bafilomycin A_1_, S-diphenylcarbazone and sodium orthovanadate, were purchased from the Sigma Chemical Company (Saint Louis, USA). Ethanol, dimethyl sulfoxide (DMSO), mercury nitrate, and triethanolamine (TEA) were from Merck (Darmstadt, Germany). The protease inhibitor cocktail (1 mmol L^-1^ benzamidine, 5 µmol L^-1^ antipain, 5 µmol L^-1^ leupeptin, 1 µmol L^-1^ pepstatin A and 5 µmol L^-1^ phenyl-methane-sulfonyl-fluoride) was from Calbiochem (Darmstadt, Germany). Ammonium sulfate-depleted PK, LDH suspensions and stock solutions of ATP, bafilomycin A_1_ and sodium orthovanadate were prepared according to Lucena et al. (2012). When necessary, enzyme solutions were concentrated on YM-10 Amicon Ultra filters. All cations were used as chloride salts.

### 2.2. Crab collections

Adult male and non-ovigerous female *U. cordatus* measuring 8-9 cm in carapace width were caught by hand from the Barra Seca mangrove (23° 24’ 58.9” S, 45° 03’ 02.9” W) in Ubatuba, São Paulo State, Brazil, during four collections made between 2015 and 2016, under ICMBio/MMA permit #29594-9 to JCM. The crabs were transported individually to the laboratory in transparent, closed plastic boxes (20×20×20 cm) containing a 3-cm deep layer of brackish water from the collection site. Before salinity acclimation in the laboratory, the crabs were maintained in their boxes for 24 h at 26 ‰S (g L^-1^, salinity) and ≈25 °C, under a natural photoperiod of 14 h light: 10 h dark.

### 2.3. Experimental design and salinity acclimation

For each salinity tested, six to eight intermolt crabs were acclimated individually to either 2, 8, 18, 26 or 35 ‰S in transparent closed plastic boxes (20×20×20 cm) containing a 3-cm deep layer of experimental medium, for 10 days at ≈25 °C, under a natural photoperiod of 14 h light: 10 h dark. The reference salinity was 26 ‰S. Salinity was adjusted by the addition of Tropic Marin sea salt to chlorine-free tap water (<0.5 ‰S). Salinities were checked daily during the acclimation period using an Atago refractometer (Warszawa, Poland). The experimental media were changed daily during the experiments, and the crabs were fed on alternate days with pieces of shrimp or fish. Uneaten food fragments were removed the following morning.

### 2.4. Preparation of the gill microsomal fraction

For each microsomal preparation, six to eight crabs were anesthetized by chilling in crushed ice for 5 min and then killed by quickly transecting the ventral ganglion with scissors and removing the carapace. The three posterior gill pairs (≈0.75 g wet weight) were rapidly excised and placed in 80 mL ice-cold homogenization buffer (20 mmol L^-1^ imidazole buffer, pH 6.8, containing 250 mmol L^-1^ sucrose, 6 mmol L^-1^ EDTA and the protease inhibitor cocktail (Lucena et al., 2012). The gills were rapidly diced and homogenized in a Potter homogenizer (600 rpm) in the homogenization buffer (20 mL buffer/g wet tissue). After centrifuging the crude extract at 20,000 ×*g* for 35 min at 4 °C, the supernatant was placed on crushed ice and the pellet was resuspended in an equal volume of homogenization buffer. After further centrifugation as above, the two supernatants were gently pooled and centrifuged at 100,000 ×*g* for 90 min at 4 °C. The resulting pellet containing the microsomal fraction was homogenized in buffer and 0.5-mL aliquots were rapidly frozen in liquid nitrogen and stored at −20 °C. No appreciable loss of (Na^+^, K^+^)-ATPase activity was seen after four-month’s storage of the microsomal preparation. All experiments were performed using gill microsomal aliquots previously incubated with alamethicin (20 μg/mg protein) for 10 min at 25 °C. Thawed aliquots were held in a crushed ice bath for no longer than 4 h.

### 2.5. Measurement of protein concentration

Protein concentration was estimated according to Read and Northcote (1981) using bovine serum albumin as the standard.

### 2.6. Continuous-density sucrose gradient centrifugation

An aliquot containing ≈3.5 mg protein of the microsomal preparation from crabs acclimated to 2, 8, 16, 25 or 35 ‰S was layered into a 10-50 % (w/v) continuous sucrose density gradient and centrifuged at 180,000 ×*g* and 4 °C for 3 h, using a Hitachi PV50T2 vertical rotor. Fractions (0.5 mL) were collected from the bottom of the gradient and were analyzed for (Na^+^, K^+^)-ATPase activity and sucrose concentration.

### 2.7. Measurement of hemolymph osmolality and Na^**+**^ **and Cl**^**-**^ concentrations

Hemolymph samples from salinity-acclimated crabs were drawn through the arthrodial membrane of the last pereiopod with a #25-7 needle coupled to an insulin syringe, frozen and held at −20 °C until analysis. Hemolymph osmolality was measured in undiluted 10-μL aliquots using a vapor pressure micro-osmometer (Model 5500, Wescor Inc., USA). Na^+^ concentration was measured after diluting the hemolymph samples by 1: 1,000 in 1% (v/v) HNO_3_, using an atomic absorption spectrophotometer (Shimadzu A-680). Chloride concentration was estimated in 10-μL aliquots by titration against mercury nitrate, employing S-diphenylcarbazone as an indicator, using a microtitrator (Model E485, Metrohm AG, Switzerland) (Santos and McNamara, 1996).

### 2.8. Measurement of (Na^**+**^, **K**^**+**^)-ATPase activity

Total ATPase activity was assayed at 25 °C using a PK/LDH coupling system in which ATP hydrolysis was coupled to NADH oxidation (Leone et al., 2015). The oxidation of NADH was monitored at 340 nm (ε_340nm, pH 7.5_= 6200 M^-1^ cm^-1^) in a Shimadzu UV-1800 spectrophotometer equipped with thermostatted cell holders. Standard conditions were 50 mmol L^-1^ HEPES buffer (pH 7.5) containing 1 mmol L^-1^ ATP (for 8 and 26 ‰S) or 0.5 mmol L^-1^ (for 2, 18 and 35 ‰S), 3 mmol L^-1^ MgCl_2_ (for 26‰S) or 2 mmol L^-1^ (for 2 and 8‰S) or 1 mmol L^-1^ (for 18 and 35 ‰S), 50 mmol L^-1^ NaCl (for 2 and 26‰S) or 30 mmol L^-1^ for 35‰S or 20 mmol L^-1^ (for 8 and 18‰S), 10 mmol L^-1^ KCl (for 2, 18 and 35‰S) or 20 mmol L^-1^ for (8 and 26‰S), 0.21mmol L^-1^ NADH 3.18 mmol L^-1^ PEP, 82 μg PK (49 U), 110 μg LDH (94 U), plus the microsomal preparation (10-30 μL), in a final volume of 1 mL. ATP hydrolysis also was estimated with 3 mmol L^-1^ ouabain; the difference in activity measured without (total ATPase activity) or with ouabain (ouabain-insensitive ATPase activity) was considered to represent the (Na^+^, K^+^)-ATPase activity.

Controls without added enzyme were also included in each experiment to quantify non-enzymatic substrate hydrolysis. Initial velocities were constant for at least 15 min provided that less than 5% of the total NADH was oxidized. Neither NADH, PEP, LDH nor PK was rate-limiting over the initial course of the assay, and no activity could be measured in the absence of NADH. (Na^+^, K^+^)-ATPase activity was checked for linearity between 10-50 μg total protein; total microsomal protein added to the cuvette always fell well within the linear range of the assay. For each ATP concentration, reaction rate was estimated in duplicate using identical aliquots from the same preparation. The mean values from the duplicates were used to fit the corresponding saturation curves, each of which was repeated three times using a different microsomal homogenate (N= 3).

### 2.9. Synthesis of [γ-^**32**^P]ATP

Synthesis of [γ-^32^P]ATP was performed as described by Walseth and Johnson (1979) as modified by Maia et al. (1988).

### 2.10. Extraction of pig kidney FXYD2 peptide

Pig kidneys were obtained from a local abattoir and the outer medullas were dissected, homogenized and the purified (Na^+^, K^+^)-ATPase was prepared according to Fontes et al. (1999). The FXYD2 peptide was then prepared according to Cortes et al. (2006). Briefly, aliquots (≈1 mg protein) of purified (Na^+^, K^+^)-ATPase were diluted 16-fold at room temperature with a methanol (46%): chloroform (46%): ammonium bicarbonate (8%) mixture (v/v) adjusted to pH 7.5. The resulting suspension was centrifuged at 1,000 *×g* for 1 min and the FXYD2-rich supernatant was dried at 40 °C in a heat block under a nitrogen stream. The dry residue was suspended in 300 μL of 50 mmol L^-1^ HEPES buffer, pH 7.5, and the suspension was used to evaluate the effect of FXYD2 on (Na^+^, K^+^)-ATPase activity.

### 2.11. Effect of FXYD2 peptide on gill (Na^**+**^, **K**^**+**^)-ATPase activity in salinity acclimated crabs

The effect of FXYD2 peptide on gill (Na^+^, K^+^)-ATPase activity of crabs acclimated to the different salinities was assayed by measuring the release of ^32^Pi from [γ-^32^P]ATP as described by Grubmeyer and Penefsky (1981) and Fontes et al. (1999). Before the reaction, aliquots containing 5 μg of gill microsomal preparation (see section 2.4) from crabs acclimated to 2, 26 or 35 ‰S were incubated with 30 μL FXYD2 peptide suspension prepared as above (1: 40 enzyme to FXYD2 ratio, v/v) at 25 °C. ATPase activity was estimated in 50 mmol L^-1^ HEPES buffer (pH 7.5) under the same ionic conditions given above (see section 2.8) in a final volume of 0.5 mL. The reaction was started by adding 2 mmol L^-1^ ATP/[γ-^32^P]ATP (specific activity 1,500 cpm/nmol). After 60 min at 25 °C, the reaction was stopped with 0.2 mL 0.4 M perchloric acid and the samples were held in a crushed ice bath. After adding 400 μL of activated charcoal, the samples were centrifuged at 700 *×g* for 5 min, and 0.5 mL aliquots (N=3) of the supernatant were collected and spotted onto a Whatman filter paper disk. The filter was dried and the ^32^Pi released was quantified by liquid scintillation counting in a Packard Tri-Carb 2100 LSC scintillation counter. Controls with acid-denatured enzyme were included in each experiment to quantify non-enzymatic substrate hydrolysis. All measurements were performed both without and with 3 mmol L^-1^ ouabain, the difference in activities being assumed to correspond to the (Na^+^, K^+^)-ATPase activity.

### 2.12. Effect of phosphorylation by endogenous protein kinases on gill microsomal (Na^**+**^, **K**^**+**^)-ATPase activity

Alamethicin-treated aliquots (see section 2.4) containing 100 µg protein of the gill microsomal (Na^+^, K^+^)-ATPase of crabs acclimated to the different salinities were assayed for phosphorylation by endogenous PKA, PKC and CaMK during 60 min. The reaction was started by adding 3 mM ATP and allowed to proceed for 60 min at 25 °C in 20 mmol L^-1^ HEPES buffer (pH 7.5), 10 mmol L^-1^ MgCl_2_, 100 mmol L^-1^ KCl, 1 mmol L^-1^ EGTA and 1 mmol L^-1^ DTT in a final volume of 0.5 mL. For PKA, the phosphorylation reaction media also contained 0.05% Triton X-100, 2.5 mmol L^-1^ dibutyryl cAMP and 3.5 μmol L^-1^ chelerythrine (PKC inhibitor). For PKC, 10 mmol L^-1^ CaCl_2_, 80 μg/μL phosphatidylserine, 100 nmol L^-1^ PMA (PKC stimulator) and 200 nmol L^-1^ H-89 were added to the reaction media. For CaMK, phosphorylation was performed by adding 10 mmol L^-1^ CaCl_2_ and 200 μg/μL calmodulin to the reaction media. Controls were also performed as above with 200 nmol L^-1^ H-89 (PKA inhibitor), 3.5 μmol L^-1^ chelerythrine (PKC inhibitor) and 2 μmol L^-1^ KN62 (CaMK inhibitor), respectively.

Aliquots (N= 3) containing 20 µg protein of protein kinase-phosphorylated gill (Na^+^, K^+^)-ATPase were then assayed for ATPase activity in 50 mmol L^-1^ HEPES buffer (pH 7.5) under the same ionic conditions given above (see section 2.8) in a final volume of 0.5 mL. The reaction was started by adding 2 mM ATP/[γ-^32^P]ATP (specific activity 1,500 cpm/nmol) and allowed to proceed for a further 60 min, at 25 °C. The reaction was stopped by adding 0.2 mL 0.4 M perchloric acid and the samples were placed in a crushed ice bath. After adding 0.4 mL activated charcoal, the samples were centrifuged at 700 *×g* for 5 min and 0.5 mL of supernatant was collected and spotted onto a Whatman filter paper disk. The filter was dried and the ^32^Pi released was quantified by liquid scintillation counting in a Packard Tri-Carb 2100 LSC liquid scintillation counter. Controls with acid-denatured enzyme were included in each experiment to quantify non-enzymatic substrate hydrolysis. All measurements were performed both without and with 3 mmol L^-1^ ouabain, and the difference in activities was assumed to correspond to the (Na^+^, K^+^)-ATPase activity.

### 2.13. SDS-PAGE analysis of the (Na^**+**^, **K**^**+**^)-ATPase α-subunit after phosphorylation by endogenous protein kinases

SDS-PAGE analyses of the protein kinase-phosphorylated proteins were performed according to Laemmli (1970), employing a 4% stacking gel and 15% resolution gel, under 60 mA constant current. Aliquots of the gill microsomal preparation (20 or 40 µg protein) were added to the phosphorylation reaction media (see section 2.12) and the reaction was started by adding 3 mM ATP/[γ-^32^P]ATP (specific activity 800,000 cpm/nmol). The reaction was stopped after 1 h by adding six volumes of electrophoresis buffer. The molecular markers in the gel were stained with colloidal Coomassie Blue, and after drying, the gel slab was autoradiographed for 24 h using a Cyclone Phosphor Imager apparatus (Perkin Elmer, Massachusetts). The images were produced by direct scanning using OptiQuant software and a proprietary storage phosphor screen.

### 2.14. Estimation of kinetic parameters

SigrafW software (Leone et al., 2005) was used to calculate the kinetic parameters V_M_ (maximum velocity), K_0.5_ (apparent dissociation constant), K_M_ (Michaelis–Menten constant), and n_H_ (Hill coefficient) values for ATP hydrolysis at the different acclimation salinities. The kinetic parameters furnished in the tables are calculated values and represent the mean ± SD derived from three different microsomal preparations (N= 3). SigrafW software can be obtained freely from http://portal.ffclrp.usp.br/sites/fdaleone/downloads.

### 2.15. Statistical analyses and calculations

Data for osmoregulatory parameters are given as the mean ± SEM (N). After meeting the criteria for normality of distribution and equality of variance, the data sets were analyzed using one-way (acclimation salinity) or two-way (acclimation salinity, presence of FXYD2 peptide) analyses of variance followed by the Student-Newman-Keuls multiple means comparison procedure to locate significant differences among treatments (SigmaPlot for Windows, version 11). Differences were considered significant at P= 0.05.

To evaluate osmotic and ionic regulatory capability, hemolymph osmolalities and [Na^+^] and [Cl^-^] were fitted to second order polynomial equations (Y= a_2_x^2^+a_1_x+a_0_) where the independent variable (x) was the osmolality of the external media. The isosmotic and iso- ionic points, represented by the intercepts of the fitted curves with the isosmotic/iso-ionic lines, were calculated according to Freire et al. (2003). Hyper- and hypo-regulatory capabilities were expressed numerically as the ratio of change in hemolymph osmolality, [Na^+^] or [Cl^-^] (Δ hemolymph parameter) compared to that of the acclimation salinity (Δ medium parameter), below or above the isosmotic or iso-ionic points, respectively. A ratio close to ‘0’ indicates excellent regulatory capability while values near ‘1’ reveal a lack of regulatory ability (Freire et al., 2003).

## 3. RESULTS

### 3.1. Hemolymph osmotic and ionic regulatory capability

*Ucides cordatus* was isosmotic (776±19 mOsm kg^-1^ H_2_O) after 10-days acclimation to 26 ‰S (780 mOsm kg^-1^ H_2_O), the reference salinity (Fig. 1). Salinity acclimation had no effect on hemolymph osmolality (P= 0.126). After acclimation to 2, 8, 18, 26 or 35 ‰S, hemolymph osmolalities were 692.2±49.4, 700.4±22.9, 720.2±85.1, 776.20±32.41, and 833.0±41.7 mOsm kg^-1^ H_2_O, respectively (Fig. 1). Hyper-osmoregulatory capability (Δ hemolymph osmolality/Δ external osmolality) was 0.12, while hypo-osmoregulatory capability was 0.21, both revealing excellent osmoregulatory ability.

**Figure 1.**
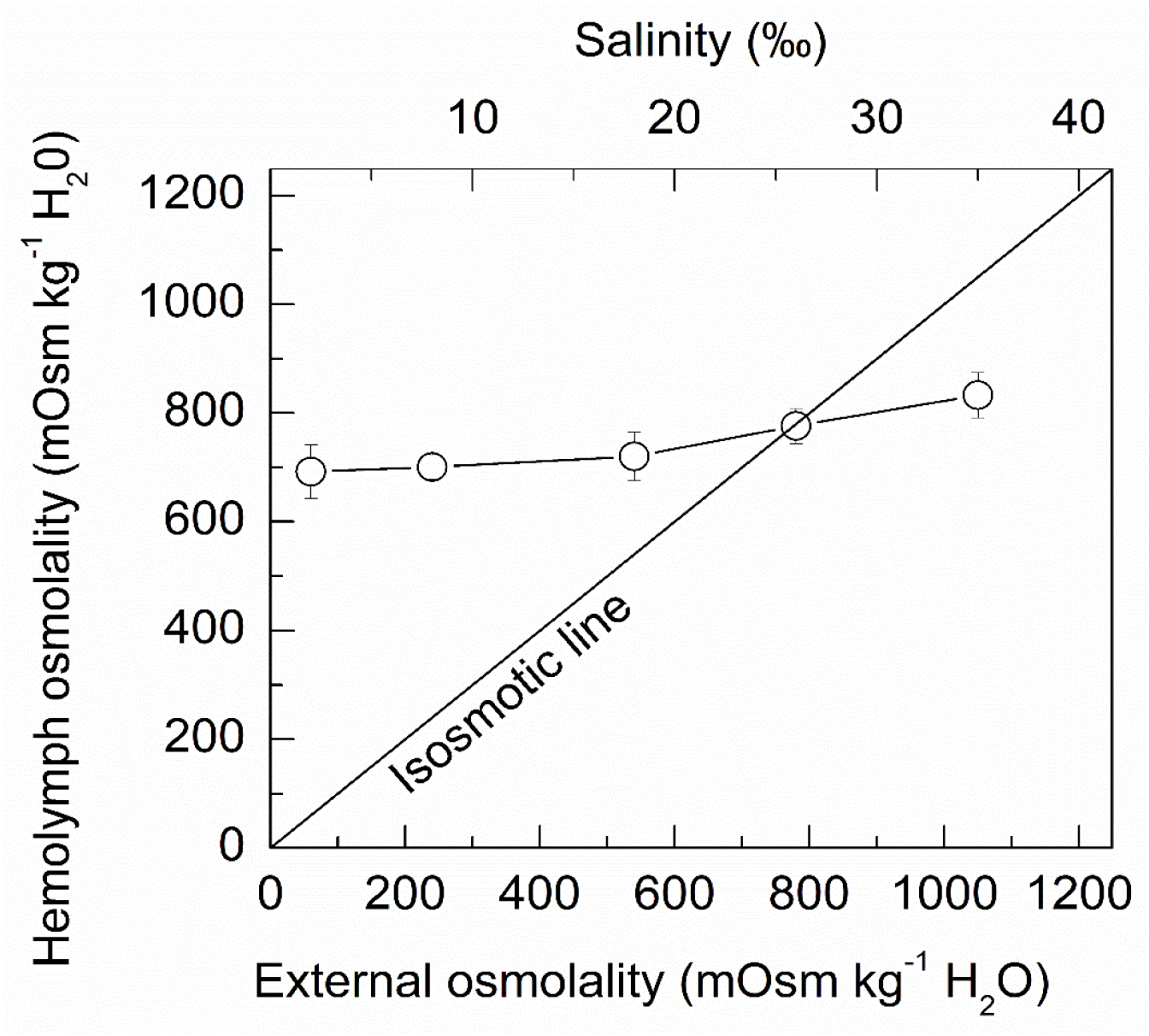
Hemolymph osmoregulatory capability in *Ucides cordatus* following hypo- or hyper-osmotic challenge for 10 days. Crabs were acclimated to 2, 8, 18, 26 (reference salinity) or 35 ‰S for 10 days. Osmolality was measured in 10-μL aliquots of hemolymph taken from individual crabs. Data are the mean ± SEM (N=5-10). The calculated isosmotic point is 776 mOsm kg^−1^ H_2_O (1 ‰S= 30 mOsm kg^−1^ H_2_O). When not visible, error bars are smaller than the symbols used.

Hemolymph chloride was iso-ionic at 22 ‰S (352 mmol L^-1^) and was hyper-regulated at 18 (364.0±29.9 mmol L^-1^), 8 (304.0±17.2 mmol L^-1^) and 2 ‰S (215.0±12.1 mmol L^-1^), but hypo-regulated at 26 (340.0±23.3 mmol L^-1^) and 35 ‰S (293.0±15.7 mmol L^-1^) (Fig. 2). Chloride hyper-regulatory ability (Δ hemolymph [Cl^-^]/Δ external [Cl^-^]) was 0.43, revealing moderate regulatory ability. Chloride hypo-regulatory ability was moderate at −0.28.

**Figure 2.**
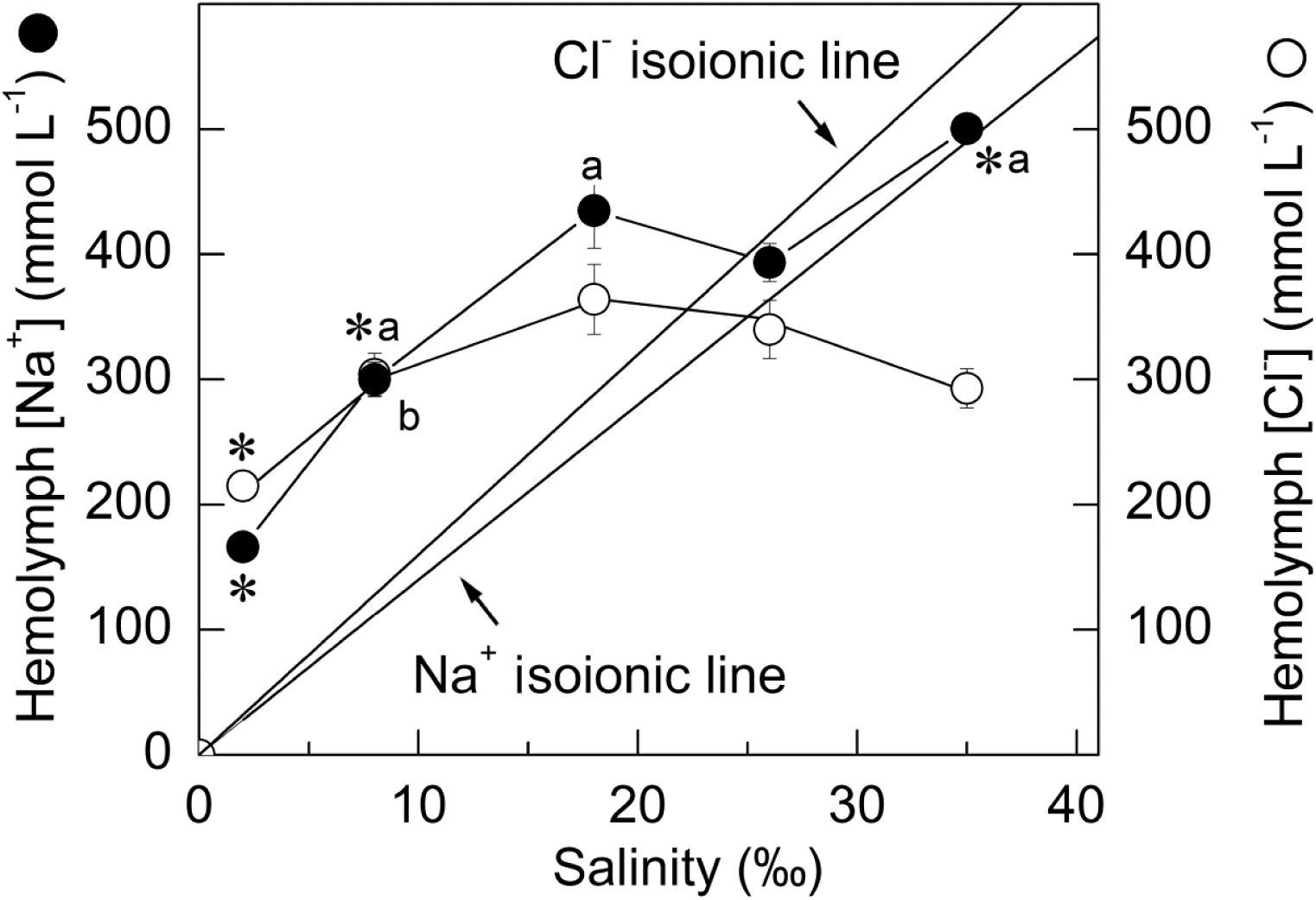
Regulation of Na^+^ and Cl^−^ concentrations in the hemolymph of *Ucides cordatus* after 10-days acclimation to different salinities. Crabs were acclimated to 2, 8, 18, 26 (reference salinity) or 35 ‰S for 10 days. Na^+^ and Cl^-^ concentrations were measured in 10-μL hemolymph aliquots taken from individual crabs. Data are the mean ± SEM (N=4-10). The isoionic points are 352 mmol L^-1^ for chloride and 490 mmol L^-1^ for sodium. 1 ‰S= 16 mmol L^-1^ Cl^-^ and 14 mmol L^-1^ Na^+^, respectively. *P≤0.05 compared to reference salinity (26 ‰S); ^a^P≤0.05 compared to immediately preceding value for sodium curve; ^b^P≤0.05 compared to immediately preceding value for chloride curve (ANOVA, SNK).

Hemolymph sodium was iso-ionic (500.4±11.0 mmol L^-1^) at 35 ‰S (490 mmol L^-1^) and hyper-regulated at all lower salinities, decreasing to 166.2±6.1 mmol L^-1^ at 2 ‰S, maintaining a ≈6:1 gradient against this medium (P< 0.001) (Fig. 2). Hemolymph Na^+^ concentrations were comparable (P= 0.108) at 18 ‰S (434.8±29.8, gradient 1.7:1) and 26 ‰S (393.6±15.2, gradient 1.1:1). Sodium hyper-regulatory ability (Δ hemolymph [Na^+^]/Δ external [Na^+^]) was weak at 0.72. Clearly, *U. cordatus* can strongly hypo- and hyper-regulate its hemolymph osmolality and Cl^-^ concentration but only weakly regulates Na^+^ concentration.

### 3.2. Effect of acclimation salinity on gill (Na^**+**^, **K**^**+**^)-ATPase activity

(Na^+^, K^+^)-ATPase activities were very different in the gill homogenates from *U. cordatus* after 10-days acclimation to the different salinities. Activity was greatest (652.4±27.1 nmol Pi min^−1^ mg^−1^ protein) in crabs acclimated to the isosmotic salinity of 26 ‰S (Fig. 3). Activities diminished by ≈50% at 18 ‰S (358.2±14.9 nmol Pi min^−1^ mg^−1^ protein) and 8 ‰S (304.9±15.2 nmol Pi min^−1^ mg^−1^ protein) and decreased markedly at 2 ‰S (24.3±1.2 nmol Pi min^−1^ mg^−1^ protein). Activity also decreased notably at 35 ‰S (45.9±2.3 nmol Pi min^−1^ mg^−1^ protein).

**Figure 3.**
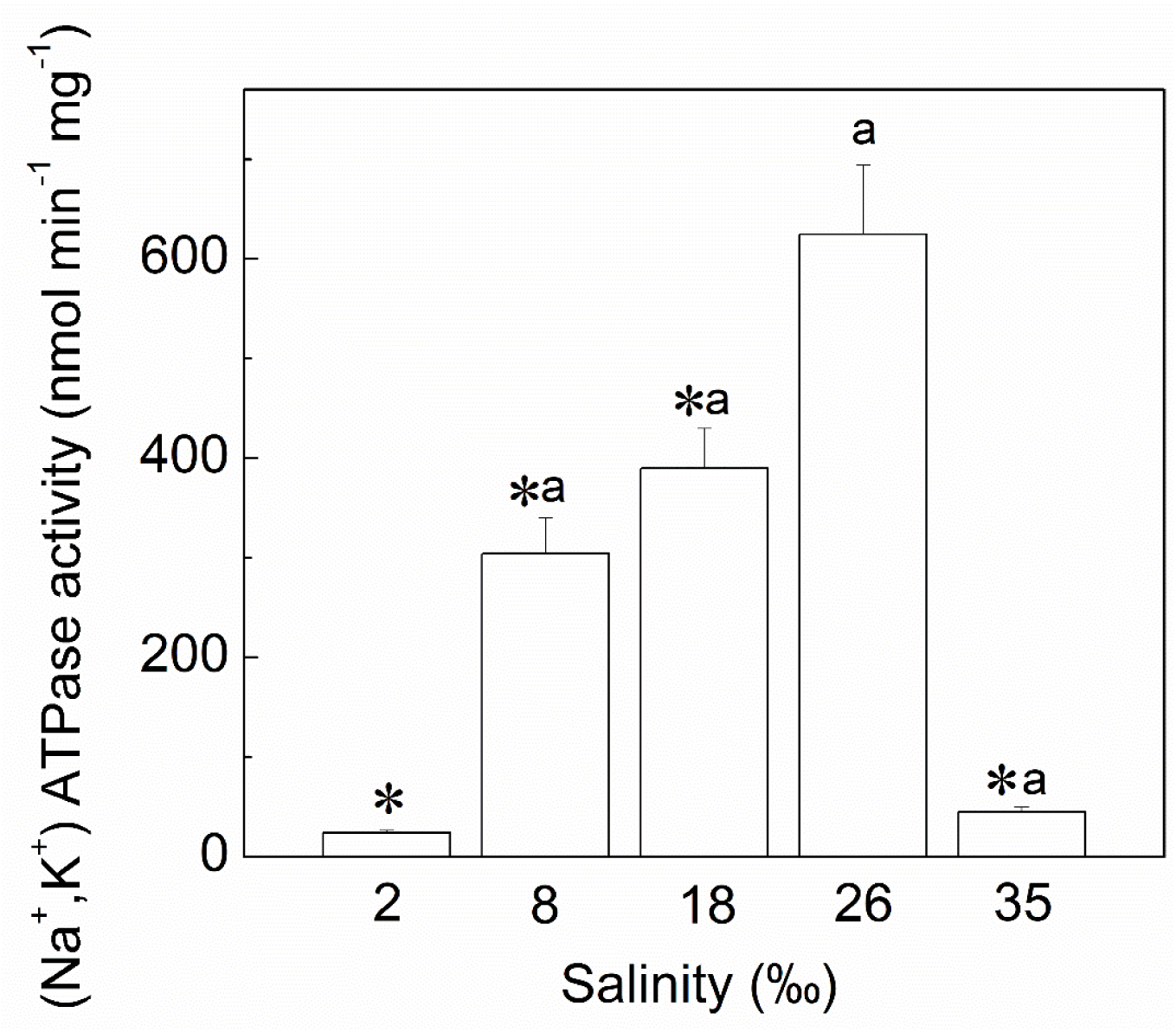
Effect of 10-days salinity acclimation on (Na^+^, K^+^)-ATPase activity in posterior gill homogenates from *Ucides cordatus*. For each salinity, activity was estimated as described in the Materials and Methods using 25, 10, 22, 20 or 28 μg protein in the assay reaction for 2, 8, 18, 26 or 35 ‰S, respectively. Mean values of duplicate measurements were used to estimate (Na^+^, K^+^)-ATPase activity at each salinity, which was repeated utilizing three different microsomal preparations (N= 3). Data are the mean±SD. *P≤0.05 compared to reference salinity (26 ‰S); ^a^P≤0.05 compared to immediately preceding value (ANOVA, SNK).

### 3.3. Effect of acclimation salinity on the modulation by ATP of gill (Na^**+**^, **K**^**+**^)-ATPase activity

Acclimation for 10 days to the different salinities markedly affected the modulation by ATP of gill (Na^+^, K^+^)-ATPase activity (Fig. 4). A single saturation curve showing Michaelis-Menten characteristics was seen over a broad range of ATP concentrations (10^−8^ to 10^−3^ mol L^−1^) for crabs acclimated at 2 ‰S, and maximum (Na^+^, K^+^)-ATPase activity was calculated as V_M_= 24.3±1.2 nmol Pi min^−1^ mg^−1^ protein and K_M_ = 29.0±2.5 μmol L^−1^ (Table 1). At 8 ‰S, a single saturation curve showing Michaelis-Menten characteristics also prevailed over the same ATP concentration range. In this case, maximum (Na^+^, K^+^)-ATPase activity was calculated as V_M_ = 304.9±15.2 nmol Pi min^−1^ mg^−1^ protein) and K_M_ = 79.1±4.7 μmol L^−1^.

**Table 1.**
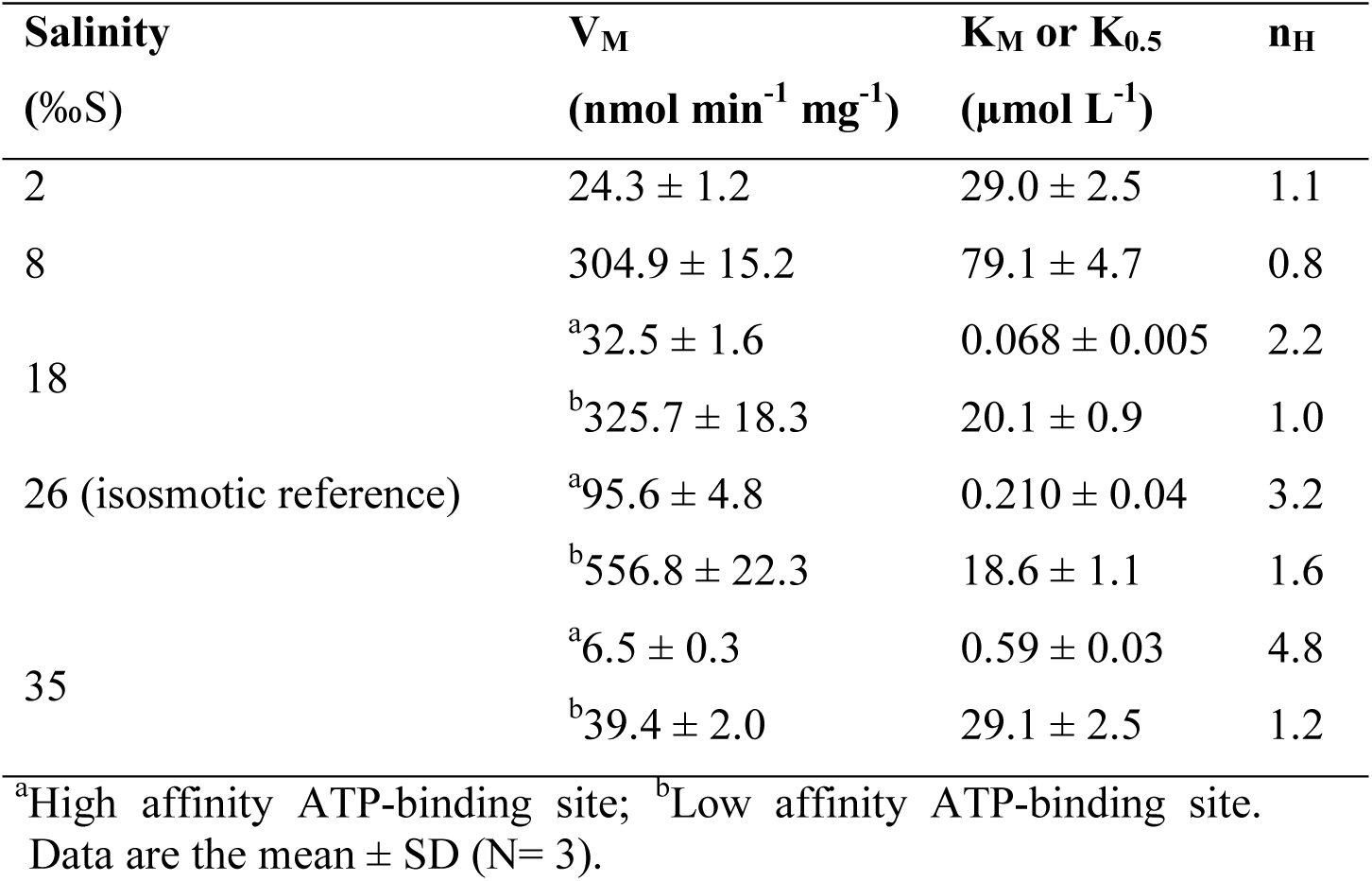
Calculated kinetic parameters for the stimulation by ATP of posterior gill (Na^+^, K^+^)-ATPase activity in *Ucides cordatus* after 10-days acclimation to different salinities.

**Figure 4.**
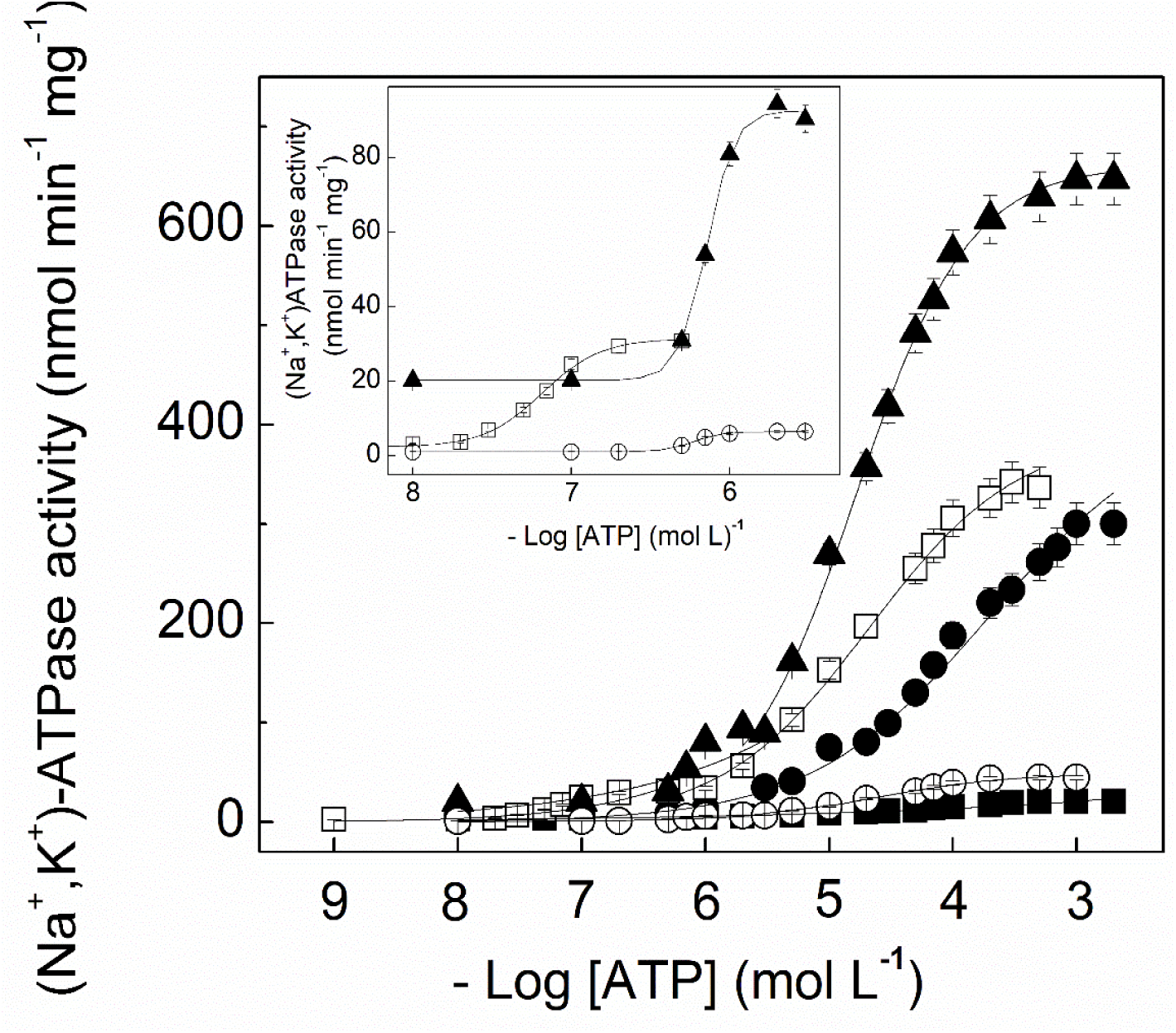
Stimulation by ATP of posterior gill (Na^+^, K^+^)-ATPase activity in *Ucides cordatus* after 10-days acclimation to different salinities. Activity was estimated as described in the Materials and Methods using 25, 10, 22, 20 or 28 μg protein in the assay reaction for 2, 8, 18, 26 or 35 ‰S, respectively. Mean values (±SD) for the duplicates were used to fit each corresponding curve, which was repeated three times using a different microsomal preparation (N= 3). (▪) 2 ‰S, (•) 8 ‰S, (□) 18 ‰S, (▴) 26 ‰S, (○) 35 ‰S. **Inset**: Effect of salinity on high affinity ATP-binding sites, (□) 18 ‰S, (▴) 26 ‰S, (○) 35 ‰S.

Acclimation to 18, 26 (reference salinity) and 35 ‰S resulted in more complex ATP saturation curves showing high (appearing at low ATP concentrations) and low affinity (appearing at high ATP concentrations) ATP-binding sites over the same ATP concentration range (inset to Fig. 4). Independently of salinity, the high affinity ATP sites showed cooperative kinetics with calculated K_0.5_ values of 0.068±0.005, 0.210±0.04 and 0.59±0.03 μmol L^−1^, respectively. Except for crabs acclimated at 18 ‰S (K_M_= 20.1±0.9 μmol L^−1^), the low-affinity ATP sites of those acclimated at 26 (K_0.5_= 18.6±1.1 μmol L^−1^) and 35 ‰S (K_0.5_= 29.1±2.5 μmol L^−1^) showed site-site interactions. Maximum gill (Na^+^, K^+^)-ATPase activity calculated for the high affinity ATP sites of crabs acclimated at 18, 26 and 35 ‰S were V_M_= 32.5±1.6, V_M_= 95.6±4.8 and V_M_= 6.5±0.3 nmol Pi min^−1^ mg^−1^ protein, respectively. The low affinity ATP sites showed V_M_= 325.7±18.3, V_M_= 556.8±22.3 and V_M_= 39.4±2.0 and nmol Pi min^−1^ mg^−1^ protein, respectively (Table 1). For the crabs acclimated to high salinities (18 to 35 ‰S), the calculated apparent dissociation constant (K_0.5_) increased with increasing salinity.

### 3.4. Continuous-density sucrose gradient centrifugation

The distribution profiles of the gill microsomal (Na^+^, K^+^)-ATPase activities of *U. cordatus* acclimated to different salinities differed along the sucrose gradient (Fig. 5). At 2 ‰S, a broad ATPase activity peak lying between 20 and 45% sucrose, showing a heavy shoulder was seen. At 8 ‰S, two well-defined activity peaks appeared between 25 and 35% sucrose (lighter fraction), and 35 and 45% sucrose (heavier fraction). At 18 ‰S, only a single well-defined activity peak lying between 25 and 40% sucrose was seen. For the 26 and 35 ‰S-acclimated crabs, the ATPase activity peak was spread along the sucrose gradient.

**Figure 5.**
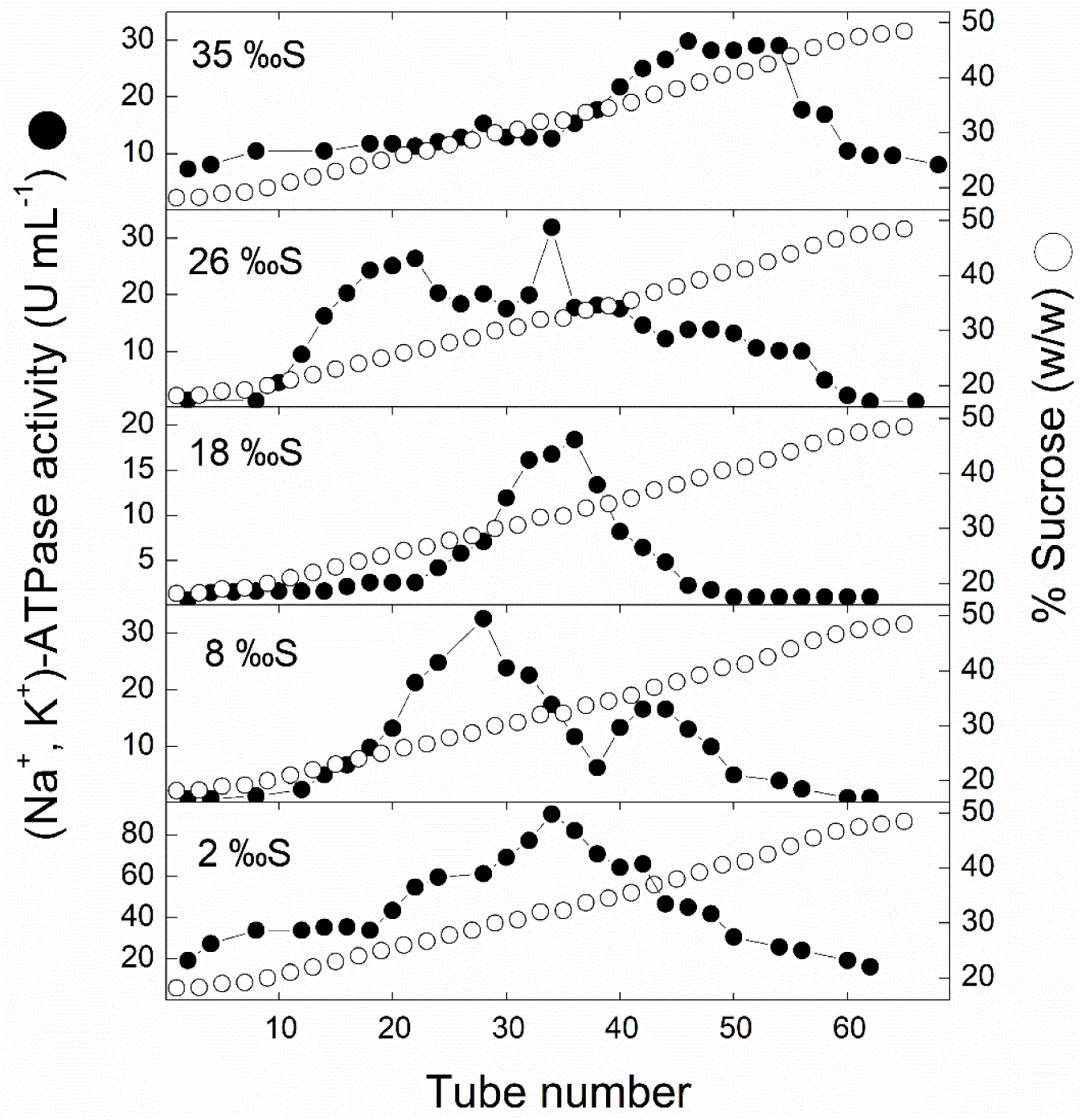
Sucrose density gradient centrifugation of (Na^+^, K^+^)-ATPase activity in a posterior gill microsomal fraction from *Ucides cordatus* after 10-days acclimation to different salinities. An aliquot containing ≈3.5 mg protein of a microsomal preparation of gill tissue from *Ucides cordatus* acclimated to each salinity was layered into a 10-50 % (w/w) continuous sucrose density gradient and centrifuged at 180,000 *×g* and 4 °C for 3 h. Fractions (0.5 mL) were collected from the bottom of the gradient and were analyzed for (Na^+^, K^+^)-ATPase activity (•) and sucrose concentration (◯).

### 3.5. Effect of exogenous FXYD2 peptide on (Na^**+**^, **K**^**+**^)-ATPase activity

(Na^+^, K^+^)-ATPase activity in the gills of crabs acclimated to 2 (hyper-osmotic condition), 26 (isosmotic, reference) or 35 ‰S (hypo-osmotic condition) for 10 days was stimulated differentially by the exogenous pig kidney FXYD2 peptide (Fig. 6 and Table 2). In the presence of FXYD2 peptide, (Na^+^, K^+^)-ATPase activity of *U. cordatus* acclimated to 2, 26 and 35 ‰S was stimulated 81, 22 and 30%, respectively. Compared to the isosmotic reference crabs, the (Na^+^, K^+^)-ATPase activity of hyperosmotic crabs was 16-fold lower while that of hypoosmotic ones was only 10-fold lower.

**Table 2.** Effect of exogenous pig kidney FXYD2 peptide on posterior gill (Na^+^, K^+^)- ATPase activity in *Ucides cordatus* after 10-days acclimation to different salinities.

| Salinity (‰S) | (Na<sup>+</sup>, K<sup>+</sup>)-ATPase activity (nmol min<sup>-1</sup> mg<sup>-1</sup> protein) |  |  |
| --- | --- | --- | --- |
|  | Control* | +FXYD2 | Stimulation (%) |
| 2 | 25.9 ± 3.5 | 47.0 ± 2.4 | 81.4 |
| 26 (isosmotic reference) | 626.3 ± 31.0 | 767.4 ± 18.0 | 22.5 |
| 35 | 55.2 ± 4.5 | 71.6 ± 3.9 | 30.0 |

Activity was estimated as described in the Materials and Methods using 5 μg of microsomal preparation and excess FXYD2 peptide. Data are the mean ± SD (N=3). \*(Na<sup>+</sup>, K<sup>+</sup>)-ATPase activity estimated without the FXYD2 peptide.

**Figure 6.**
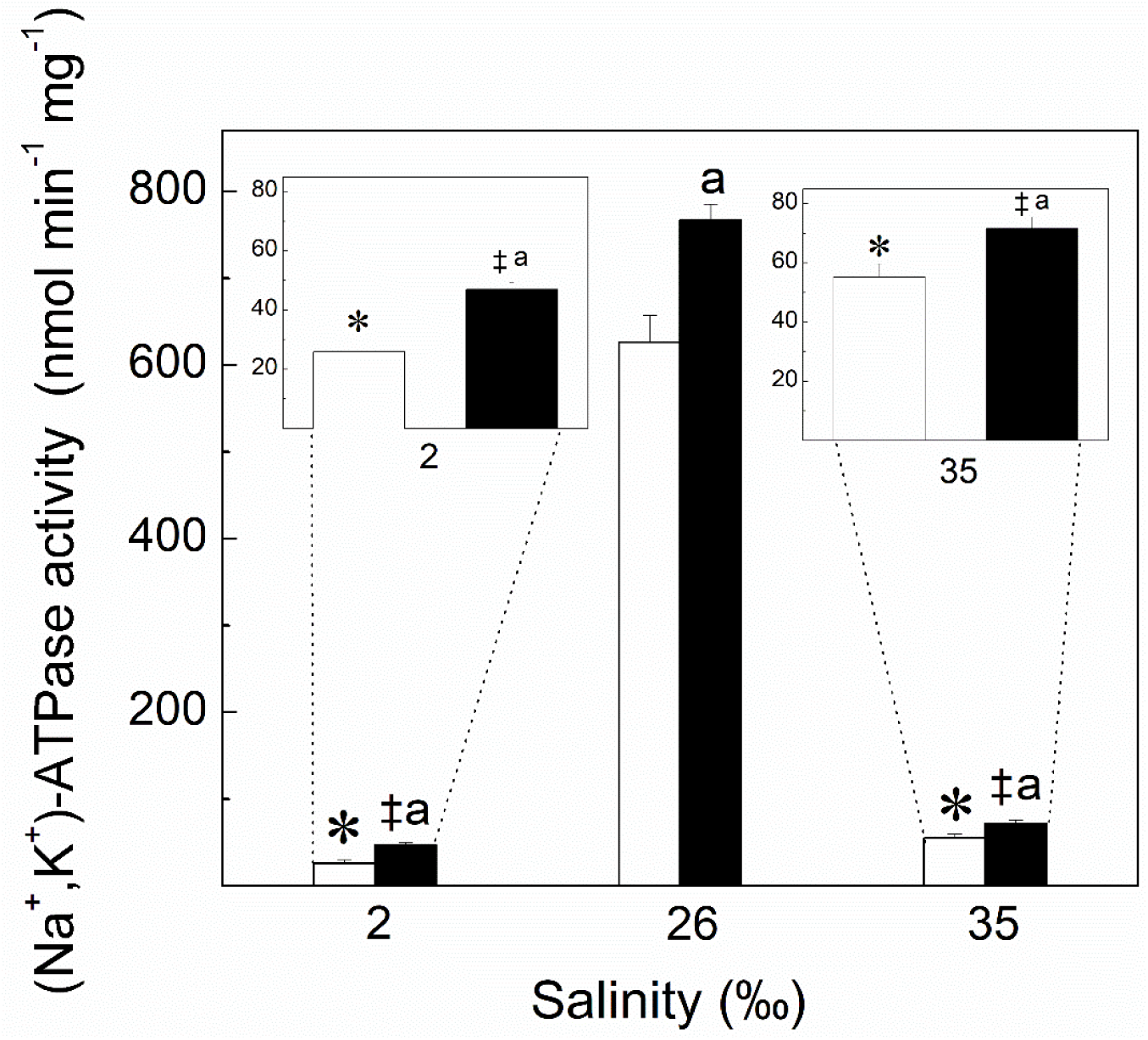
Stimulation by pig kidney FXYD2 peptide of posterior gill (Na^+^, K^+^)-ATPase activity in *Ucides cordatus* after 10-days acclimation to different salinities. (Na^+^, K^+^)-ATPase activity was estimated as described in the Materials and Methods using 5 μg protein from the microsomal fraction for each salinity and excess exogenous pig kidney FXYD2 in the assay reaction. Mean values (±SD) for the duplicates were used to estimate the (Na^+^, K^+^)-ATPase activity at each salinity, which was repeated three times using a different microsomal preparation (N= 3). *P≤0.05 and ^‡^P≤0.05 compared to reference value (26 ‰S) without or with FXYD2, respectively; ^a^P<0.05 compared to the same salinity without FXYD2 (two-way ANOVA, SNK). (□)- without FXYD2. (▪)- with FXYD2.

### 3.6. Effect of phosphorylation by endogenous protein kinases on gill (Na^**+**^, **K**^**+**^)-ATPase activity

Protein kinases A, C and CaMK all inhibited the gill (Na^+^, K^+^)-ATPase activity of *U. cordatus* acclimated to different salinities (Table 3). In the presence of dibutyryl cAMP (PKA stimulator), the activity of the isosmotic reference crabs (26 ‰S) was inhibited by ≈95% while that of 2 ‰S and 35 ‰S-acclimated crabs was inhibited by ≈50 and ≈35%, respectively. However, inhibition in the isosmotic crabs was ≈90%, and reversed by H89 (PKA inhibitor). Similarly, inhibition was fully reversed in the hyper- (2 ‰S) and hypo-osmoregulating (35 ‰S) crabs. These data demonstrate that gill (Na^+^, K^+^)-ATPase activity is regulated by PKA, independently of salinity.

**Table 3.** Effect of protein kinases A and C and calmodulin on posterior gill (Na^+^, K^+^)-ATPase activity in *Ucides cordatus* after 10-days acclimation to different salinities.

| Salinity (‰S) | (Na<sup>+</sup>, K<sup>+</sup>)-ATPase activity (nmol min<sup>-1</sup> mg<sup>-1</sup> protein) |  |  |  |  |  |  |
| --- | --- | --- | --- | --- | --- | --- | --- |
|  | Control* | PKA |  | PKC |  | CaMK |  |
|  |  | - H89 | + H89 | - Chelerythrine | + Chelerythrine | - KN62 | + KN62 |
| 2 | 24.2±1.1 | 12.1±1.0 | 22.53±0.1 | 15.9±0.2 | 19.0±0.3 | 17.5±0.3 | 18.1±1.8 |
| 26 (isosmotic reference) | 617.1±14.1 | 35.6±5.5 | 539.64±7.7 | 72.3±7.6 | 220.5±3.2 | 265.9±2.4 | 406.3±3.8 |
| 35 | 52.2±1.3 | 35.3±2.1 | 60.9±3.7 | 21.6±2.0 | 56.6±1.7 | 38.1±2.3 | 56.2±5.2 |

\*(Na<sup>+</sup>, K<sup>+</sup>)-ATPase activity estimated without protein kinase stimulators (see section 2.12).

In the presence of PMA (PKC stimulator), PKC also differentially inhibited gill (Na^+^, K^+^)-ATPase activity in the crabs acclimated to 2, 26 and 35 ‰S by ≈35%, ≈90% and ≈60%, respectively (Table 3). Chelerythrine (PKC inhibitor) notably reversed the inhibition seen in the 2 ‰S-acclimated (≈80% recovery), and completely in the 35 ‰S-acclimated crabs. However, recovery of the (Na^+^, K^+^)-ATPase activity was only 35% in the isosmotic crabs (26 ‰S). These data suggest that the gill (Na^+^, K^+^)-ATPase activity can be regulated by PKC in a salinity-dependent fashion.

Calmodulin, in the presence of calcium, also inhibited gill (Na^+^, K^+^)-ATPase activity, due to stimulation of endogenous CaMK, in a salinity-dependent fashion. Inhibition was more pronounced in the isosmotic reference crabs (≈60%) than at 2 (≈30%) and 35 ‰S (25%). Activity was completely recovered in the presence of KN62 (specific CaMK inhibitor) in the hypo-osmoregulating (35 ‰S) crabs, but was only partially reversed in the isosmotic (≈65% recovery) and hyperosmotic crabs (≈75%). These data suggest that gill (Na^+^, K^+^)-ATPase activity can be regulated by CaMK in a salinity-dependent fashion. This is the first demonstration of inhibitory phosphorylation of a crustacean (Na^+^, K^+^)-ATPase by Ca^2+^/calmodulin-dependent kinase.

### 3.7. SDS-PAGE autoradiography of (Na^**+**^, **K**^**+**^)-ATPase subunits phosphorylated by protein kinases

Phosphorylation by endogenous PKA of the α- and γ-subunits of the gill (Na^+^, K^+^)- ATPase was greatest in the 2 ‰S-acclimated crabs (Fig. 7A, lanes 1 and 2) and less intense in the isosmotic reference crabs at 26 ‰S (Fig. 7A, lanes 4 and 5) and in those at 35 ‰S (Fig. 7A, lanes 7 and 8). The PKA inhibitor H89 completely inhibited phosphorylation of these subunits in the 2 ‰S-acclimated crabs (Fig. 7A, lane 3), and to a lesser extent in the reference crabs (26 ‰S) (Fig. 7A, lane 6) and those in 35 ‰S (Fig. 7A, lane 9). Endogenous PKC also differentially phosphorylated the α-subunit in the 35 ‰S- (Fig. 7B, lanes 7 and 8) and 26 ‰S-acclimated crabs (Fig. 7B, lanes 4 and 5), and to a lesser extent in the 2 ‰S-acclimated crabs (Fig. 7B, lanes 1 and 2). Chelerythrine almost completely reversed phosphorylation of the α-subunit in the 2 ‰S-acclimated crabs (Fig. 7B, lane 3), and partially in the 26 ‰S- (Fig. 7B, lane 6) and 35 ‰S-acclimated crabs (Fig. 7B, lane 9). Stimulation by calmodulin resulted in phosphorylation of the α-subunit only in crabs acclimated to 35 ‰S (Fig. 7C, lanes 7 and 8), and to a lesser degree in the 26 ‰S-acclimated crabs (Fig. 7C, lanes 4 and 5). No phosphorylation of the α-subunit was seen in the 2 ‰S-acclimated crabs (Fig. 7C, lanes 1 and 2). KN62 completely reversed α-subunit phosphorylation in the 35 ‰S-acclimated crabs (Fig. 7C, lane 9) but only partially in the reference crabs (Fig. 7C, lane 6).

**Figure 7.**
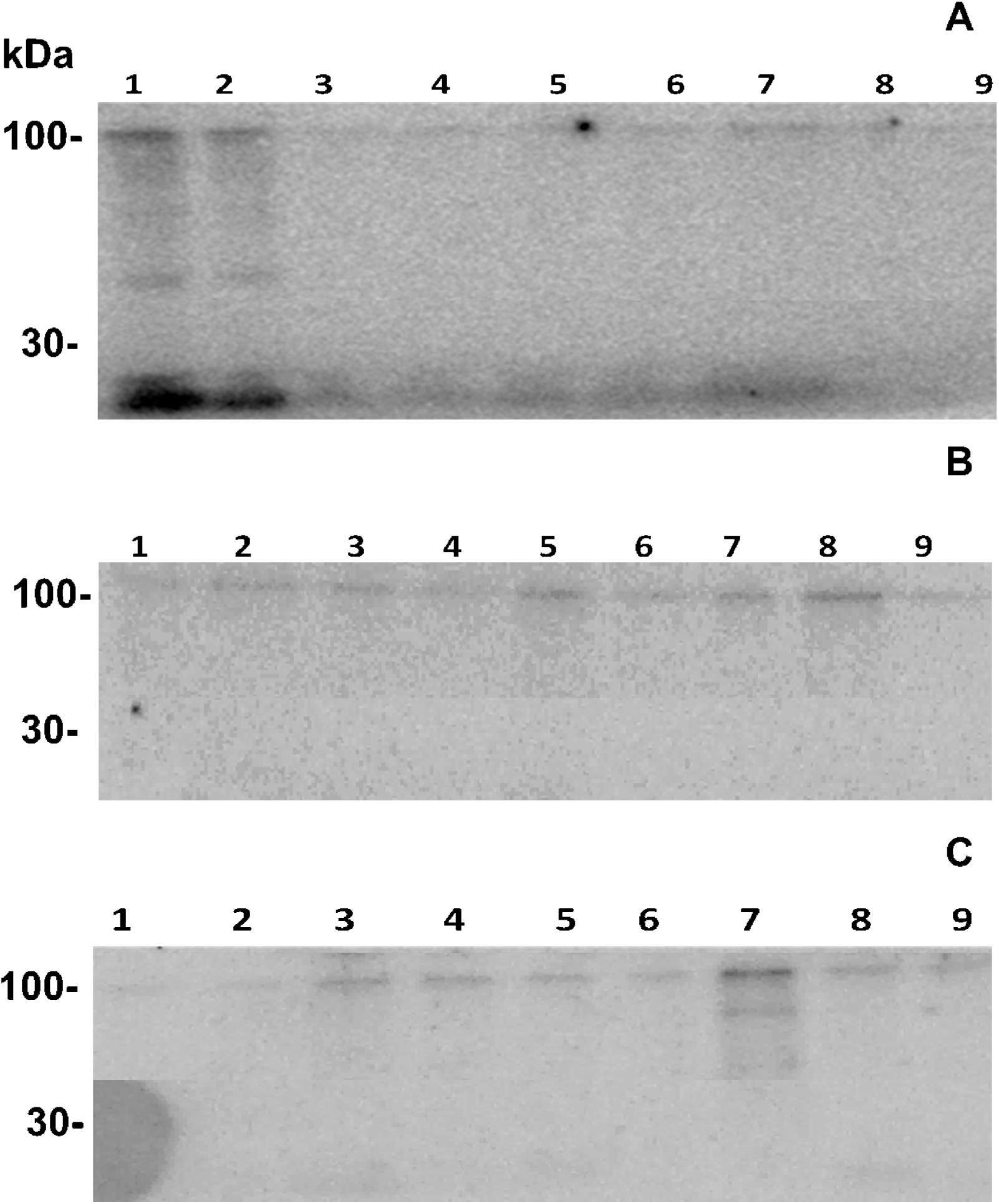
Phosphorylation of the posterior gill (Na^+^,K^+^)-ATPase by endogenous protein kinases A and C and Ca^2+^/calmodulin-dependent protein kinase. **A-** SDS-PAGE autoradiography of proteins in the gill microsomal fraction phosphorylated by endogenous PKA. Lanes 1, 4 and 7: gill microsomal fraction (20 µg protein) from crabs acclimated to 2, 26 or 35 ‰S, respectively, with 2.5 mmol L^-1^ db-cAMP, a PKA activator. Lanes 2, 5 and 8: gill microsomal fraction (40 µg protein) from crabs acclimated to 2, 26 or 35 ‰S, respectively, with 2.5 mmol L^-1^ db-cAMP. Lanes 3, 6 and 9: gill microsomal fraction (20 µg protein) from crabs acclimated to 2, 26 or 35 ‰S, respectively, with 200 nmol L^-1^ H89, a PKA inhibitor. **B-** SDS-PAGE autoradiography of proteins in the gill microsomal fraction phosphorylated by endogenous PKC. Lanes 1, 4 and 7: gill microsomal fraction (20 µg protein) from crabs acclimated to 2, 26 or 35 ‰S, respectively, with 80 μg/μL phosphatidylserine and 100 nmol L^-1^ PMA, a PKC stimulator. Lanes 2, 5 and 8: gill microsomal fraction (40 µg protein) from crabs acclimated to 2, 26 or 35 ‰S, respectively, with 80 μg/μL phosphatidylserine and 100 nmol L^-1^ PMA. Lanes 3, 6 and 9: gill microsomal fraction (20 µg protein) from crabs acclimated to 2, 26 or 35 ‰S, respectively, with 3.5 µmol L^-1^ chelerythrine, a PKC inhibitor. **C-** SDS-PAGE autoradiography of proteins in the gill microsomal fraction phosphorylated by endogenous Ca^2+^/calmodulin-dependent kinase. Lanes 1, 4 and 7: gill microsomal fraction (20 µg protein) from crabs acclimated to 2, 26 or 35 ‰S, respectively, with 100 μg/μL calmodulin. Lanes 2, 5 and 8: gill microsomal fraction (40 µg protein) from crabs acclimated to 2, 26 or 35 ‰S, respectively, with 100 μg/μL calmodulin. Lanes 3, 6 and 9: gill microsomal fraction (20 µg protein) from crabs acclimated to 2, 26 or 35 ‰S, respectively, with 100 μg/μL calmodulin and 2 µmol L^-1^ KN62, a CaMK inhibitor. Molecular weight markers (30 and 100 kDa, Magic Markers, ThermoFisher Scientific) indicated at left of panel. α-subunit, ≈100 kDa, FXYD2, ≈7 kDa).

### 3.8. Effect of acclimation salinity on P-ATPase activities in the gill microsomal preparation

Salinity acclimation altered the amount of (Na^+^, K^+^)-ATPase activity present in the microsomal preparation (Table 4). Over the salinity range employed, ouabain inhibited 50 to 85% of (Na^+^, K^+^)-ATPase activity. Systematic inhibition of the microsomal preparation using ouabain together orthovanadate disclosed considerable (≈50%), different P-ATPase activities in the ouabain-insensitive ATPase activity of crabs acclimated at 2 ‰S; neutral phosphatases constituted the main P-ATPases. Inhibition using ouabain together with ethacrynic acid showed that 50-60% of the ouabain-insensitive ATPase activity consists of Na^+^- or K^+^- stimulated ATPase in crabs acclimated at 18, 26 and 35 ‰S. High V(H^+^)- and Ca^2+^-ATPase activities were detected in the ouabain-insensitive ATPase activity of crabs acclimated at 2 ‰S.

**Table 4.** Effect of various inhibitors on total ATPase activity in a microsomal preparation from the posterior gills of *Ucides cordatus* after 10-days acclimation to different salinities.

| Condition | (Na<sup>+</sup>, K<sup>+</sup>)-ATPase activity (nmol min<sup>-1</sup> mg<sup>-1</sup> protein) |  |  |  |  | ATPase type likely present |
| --- | --- | --- | --- | --- | --- | --- |
|  | 2 ‰S | 8 ‰S | 18 ‰S | 26 ‰S* | 35 ‰S |  |
| Control | 40.3 ± 2.5 | 399.2 ± 20.0 | 412.2 ± 21.2 | 774.6 ± 2.3 | 59.3 ± 2.5 | Total ATPase |
| Ouabain (5 mmol L<sup>-1</sup>) | 18.6 ± 1.6 | 100.3 ± 5.0 | 58.5 ± 3.2 | 131.5 ± 3.4 | 14.8 ± 0.9 | (Na<sup>+</sup>, K<sup>+</sup>)- |
| Orthovanadate (50 µmol L<sup>-1</sup>) | 7.9 ± 1.0 | 102.5 ± 4.6 | 58.4 ± 2.5 | 131.0 ± 2.2 | 14.8 ± 0.8 | P-ATPase |
| Ouabain + Orthovanadate | 8.2 ± 1.0 | 100.6 ± 3.5 | 59.3 ± 3.8 | 128.5 ± 2.5 | 15.8 ± 1.0 | - |
| Ouabain + 10 µmol L<sup>-1</sup> Aurovertin | 9.2 ± 1.1 | 31.3 ± 1.5 | 46.5 ± 2.5 | 46.7 ± 2.3 | 14.7 ± 0.9 | F<sub>0</sub>F<sub>1</sub>- |
| Ouabain + 4 µmol L<sup>-1</sup> Bafilomycin | 5.3 ± 0.8 | 73.0 ± 2.9 | 42.7 ± 3.0 | 100.6 ± 3.5 | 10.5 ± 0.5 | V(H<sup>+</sup>)- |
| Ouabain + 2 mmol L<sup>-1</sup> Ethacrynic acid | 16.4 ± 1.7 | 100.1 ± 5.2 | 24.2 ± 1.0 | 131.1 ± 1.5 | 6.8 ± 0.3 | Na<sup>+</sup>- or K<sup>+</sup>- |
| Ouabain + 5 mmol L<sup>-1</sup> Theophylline | 7.2 ± 1.1 | 97.3 ± 4.3 | 55.1 ± 2.6 | 126.8 ± 1.1 | 14.7 ± 0.8 | NP* |
| Ouabain + 0.5 µmol L<sup>-1</sup> Thapsigargin | 4.2 ± 0.8 | 100.7 ± 3.8 | 55.6 ± 1.8 | 127.3 ± 0.8 | 15.2 ± 0.9 | Ca<sup>2+</sup> |
| Ouabain + 1 mmol L<sup>-1</sup> EGTA | 5.1 ± 0.6 | 99.1 ± 2.9 | 56.8 ± 3.0 | 126.5 ± 1.0 | 14.3 ± 0.8 | Ca<sup>2+</sup> |
| Ouabain + 20 µL Ethanol | 19.7 ± 2.4 | 100.8 ± 5.4 | 61.3 ± 4.9 | 130.2 ± 1.8 | 14.1 ± 0.7 | - |
| Ouabain + 20 µL DMSO | 18.6 ± 2.0 | 99.9 ± 4.7 | 62.7 ± 5.23 | 130.6 ± 0.9 | 14.7 ± 0.6 | - |

\*NP= neutral phosphatases. Data are the mean ± SD (N=3). \*26 ‰S represents the isosmotic reference salinity.

## 4. DISCUSSION

This investigation shows that the acclimation of *U. cordatus* to salinities from 2 to 35 ‰S has a negligible effect on hemolymph osmolality, which is strongly hyper- and hypo-regulated. Hemolymph Cl^-^ and Na^+^ are less well regulated. At salinities above 18‰S, the posterior gill (Na^+^, K^+^)-ATPase exhibits an additional high-affinity ATP binding site that corresponds to 10-20% of the total activity. (Na^+^, K^+^)-ATPase activity is stimulated by exogenous FXYD2 peptide but phosphorylation by PKA, PKC and CaMK inhibits activity. The inhibition by CaMK is the first report of regulatory phosphorylation of the crustacean gill (Na^+^, K^+^)-ATPase.

The different profiles of (Na^+^, K^+^)-ATPase activity revealed by sucrose gradient centrifugation may derive from their origin in membrane fragments with distinct lipid to protein ratios induced by salinity acclimation (Furriel et al., 2010; Lucena et al., 2012). The lipid environment affects (Na^+^, K^+^)-ATPase and other P_II_-type ATPase activities such as the Ca^2+^-ATPase through physico-chemical interactions (Cornelius et al., 2015), and membrane lipid composition may affect membrane permeability influencing ion and water fluxes at low salinity (Long et al., 2019).

On osmotic challenge by acclimation to different salinities, *U. cordatus* strongly hyper-/hypo-regulates hemolymph osmolality and [Cl^-^], with [Na^+^] being less regulated, revealing independent adjustment of these ions, as seen in other crustaceans (Freire et al., 2003; Kirschner, 2004; Faleiros et al., 2010). The crab’s osmoregulatory abilities appear to sustain the use of a wide variety of habitats, including mangrove forests and intertidal areas. Some terrestrial and semi-terrestrial species like *Cardisoma carnifex* resist lengthy desiccation, showing only small changes in hemolymph Na^+^ (Wood et al., 1986). Semi-terrestrial crabs such as *U. cordatus* (Harris and Santos, 1993a), *Birgus latro* (Morris et al., 1991), *Gecarcinus lateralis* and *Ocypode quadrata* (Wolcott and Wolcott, 1985) can reprocess urine in their gill chambers reabsorbing urinary excreted salt across the gill epithelia. This ability likely reflects physiological adaptation to an environment in which salinity variation and periodic or complete emersion are frequent.

As crustacean hemolymph become isosmotic and iso-natremic at high salinities, mRNA transcription of the gill (Na^+^, K^+^)-ATPase α-subunit is down regulated, resulting in reduced enzyme expression and activity (Luquet et al., 2005; Faleiros et al., 2018). The diminished gill (Na^+^, K^+^)-ATPase activity seen in *U. cordatus* at 35 ‰S is consistent with this finding and may account for the crab’s inability to excrete Na^+^, which is iso-ionic. Hemolymph Cl^-^, however, is strongly hypo-regulated, evidently by a mechanism less dependent on the gill (Na^+^, K^+^)-ATPase such as the sodium-potassium two-chloride symporter (McNamara and Faria, 2012) or a Cl^-^-stimulated ATPase (Gerencser, 1996). However, despite studies suggesting the participation of a Cl^-^-stimulated ATPase, together with anion-coupled antiports and sodium-coupled symports, in the same membrane system, there is no direct evidence for primary, active Cl^-^ transport (for review see Gerencser, 1996).

Gill (Na^+^, K^+^)-ATPase activity also decreased progressively and markedly with acclimation to lower salinities (18, 8 and 2 ‰S, see Fig. 3), which is unusual, since activities generally increase at low salinities, counterbalancing passive Na^+^ efflux (Lucu and Towle, 2003; Luquet et al., 2005; Garçon et al., 2009; Antunes et al., 2017; Faleiros et al., 2018). This decrease may derive from our use of crabs acclimated under emerged rather than submerged conditions, with free access to their experimental media. Reprocessing of a largely isosmotic and iso-ionic urine by the gills in emerged crabs may be less demanding energetically than is ion uptake from hypo-osmotic media in submerged crabs (*i. e*., against a 6:1 Na^+^ gradient in 2 ‰S) requiring less (Na^+^, K^+^)-ATPase based transport activity. Hyper-osmoregulation in *U. cordatus* is clearly driven by a mechanism not primarily dependent on the (Na^+^, K^+^)-ATPase. Hemolymph Na^+^ and Cl^-^ uptake in dilute media may be maintained by ion transporters like the Na^+^/H^+^ and Cl^-^/HCO_3_^-^ antiporters, the Na^+^/K^+^/2Cl^-^ symporter, and a V(H^+^)-ATPase/apical Na^+^ channel arrangement (Kirschner, 2005; Genovese et al., 2005; Freire et al., 2008; McNamara and Faria, 2012).

The osmolality of crustacean hemolymph depends mainly on its Na^+^ and Cl^-^ concentrations (Péqueux, 1995). In *U. cordatus* at 26 ‰S, hemolymph [Na^+^] (≈390 mmol L^-1^) and [Cl^-^] (≈340 mmol L^-1^) account for ≈94% of osmolality (≈780 mOsm kg^-1^ H_2_O). At 2 ‰S, [Na^+^] (≈170 mmol L^-1^) and [Cl^-^] (≈220 mmol L^-1^) contribute just 56% (≈692 mOsm kg^-1^ H_2_O) to osmolality (see Figs. 1 and 2). Osmolytes other than Na^+^ and Cl^-^, such as free amino acids (Augusto et al., 2007) and NH_4_^+^, may sustain the elevated hemolymph osmolality in dilute media, further reducing (Na^+^, K^+^)-ATPase activity and dependence on Na^+^. Antennal gland (Na^+^, K^+^)-ATPase activity increases in *U. cordatus* acclimated to low salinity (Harris and Santos, 1993b), suggesting augmented ion reabsorption from the urine.

SDS-PAGE autoradiography confirmed the phosphorylation by PKA of the (Na^+^, K^+^)- ATPase α- and γ-subunits in hyper-osmoregulating crabs (see Fig. 7A, lanes 1 and 2).

However, phosphorylation of the γ-subunit does not appear to contribute to overall (Na^+^, K^+^)-ATPase activity. Despite the ≈80% increase in the presence of exogenous FXYD2 (see Table 2), this activity represents only ≈6% of (Na^+^, K^+^)-ATPase activity compared to isosmotic reference crabs (Fig. 6). PKA-induced inhibition was almost completely reverted by the inhibitor H89, suggesting that PKA may phosphorylate α-subunit Ser_943_ as seen in rat kidney COS cells (Cheng et al., 1997). The phosphorylation of the *U. cordatus* gill microsomal (Na^+^, K^+^)-ATPase by endogenous CaMK is a novel finding and is the first demonstration of inhibitory phosphorylation of a crustacean (Na^+^, K^+^)-ATPase by a Ca^2+^/calmodulin-dependent kinase. The differential abilities of the various kinases to phosphorylate their targets may derive from their expression levels, which may diverge under different salinity conditions, or from the availability of endogenous modulators, as seen in *Chasmagnathus granulata* (Halperin et al., 2004), *Callinectes sapidus* (Arnaldo et al., 2014) and *Litopenaeus vannamei* (Xu et al., 2016).

(Na^+^, K^+^)-ATPase kinetic behavior was altered as a function of acclimation salinity (Fig. 4). Crabs acclimated to 2 and 8 ‰S exhibited typical Michaelis-Menten behavior, K_M_ increasing ≈3-fold and V_M_ ≈15-fold in the latter salinity (Table 1). For 18-, 26- and 35 ‰S- acclimated crabs, in addition to the exposure of a high affinity ATP-binding site, the enzyme also showed allosteric behavior (Fig. 4 and inset). While the K_0.5_ of the low affinity ATP-binding site was unaltered with increasing acclimation salinity, binding by the high-affinity site increased ≈10-fold (Table 1). (Na^+^, K^+^)-ATPase isoforms showing high and low affinity ATP-binding sites are present in many crustacean gill epithelia (Masui et al., 2002; Lucu and Towle, 2003; Leone et al., 2017; Farias et al., 2017). The non-exposure of the high affinity site after acclimation of *U. cordatus* to dilute media is similar to findings for the hermit crab *C. symmetricus* (Faleiros et al., 2018; and Antunes et al., 2017 as *C. vittatus*), the blue crab *Callinectes danae* (Masui et al., 2009) and the rock crab *Cancer pagurus* (Gache et al., 1976). ATP is considered to play both a catalytic and an allosteric role in the (Na^+^, K^+^)-ATPase reaction cycle (Beaugé et al., 1997; Krumscheid et al., 2004), and high and low affinity ATP-binding sites on the (Na^+^, K^+^)-ATPase are well known (Glynn, 1985; Ward and Cavieres, 1998). However, despite the plethora of crystallographic data suggesting a binding site within the (Na^+^, K^+^)-ATPase N-domain (Kanai et al., 2013; Nyblon et al., 2013; Chourasia and Sastry, 2012; Shinoda et al., 2009; Morth et al., 2007), its exact localization on the enzyme molecule remains an open question (Krumscheid et al., 2004; Morth et al., 2007). Regulatory phosphorylation by protein kinases can alter the kinetic profile of the host enzyme, *e. g*., phosphorylation by PKA of liver phosphofructokinase II dramatically changes kinetic behavior in response to glucagon (Pilkis et al., 1988, 1995).

Small amphipathic peptides that carry the FXYD motif such as the FXYD1 to FXYD12 series are known to bind to and directly regulate P-type ATPases (Geering, 2006; Arystarkhova et al., 2007). The blue crab *Callinectes danae* was the first crustacean shown to express the FXYD2 subunit, a 6.5-kDa protein recognized by a γC33 polyclonal anti-FXYD2 antibody, phosphorylated by endogenous PKA (Silva et al., 2012). Phosphorylated pig kidney FXYD2 stimulates the *C. danae* gill (Na^+^, K^+^)-ATPase by ≈40% (Silva et al., 2012). Similarly, the gill (Na^+^, K^+^)-ATPase of 2-, 26- and 35 ‰S-acclimated *U. cordatus* also is stimulated by exogenous phosphorylated pig kidney FXYD2 peptide, being greater in hyper- (≈80%) than in hypo-osmoregulating crabs (≈30%) (Fig. 6, Table 3). The ≈50% activation by exogenous FXYD2 seen at both low and high salinities in *U. cordatus* (Fig. 6) is comparable to that for *C. danae* (Silva et al., 2012). Phosphorylation of endogenous FXYD2 peptide by endogenous PKA in *U. cordatus* was greatest at 2 ‰S (see Fig. 7) as seen in gills of the diadromous salmon *Salmo salar* (Tipsmark, 2008), euryhaline pufferfish *Tetraodon nigroviridis* (Wang et al., 2008), and the euryhaline milkfish *Chanos chanos* (Yang et al., 2019a), which express the FXYD11 isoform. While interaction of the FXYD11 peptide with the (Na^+^, K^+^)-ATPase has been intensively investigated in fish gills (Tipsmark et al., 2010; Yang et al., 2013; Chang et al., 2016; Liang et al., 2017; Yang et al., 2019a), information on the functional interaction of the FXYD2 peptide with the (Na^+^, K^+^)-ATPase is scant (Silva et al., 2012; Yang et al., 2019b). While the present study has revealed regulatory effects of the FXYD2 peptide, its role in the physiological acclimation of crustaceans to different salinities remains unclear.

Salinity acclimation affects not only (Na^+^, K^+^)-ATPase activity but also that of various ouabain-insensitive ATPases in the gill microsomal preparation (Table 4). Most activities decrease at low and high acclimation salinities compared to the isosmotic crabs. *Ucides cordatus* clearly exhibits a complex assemblage of osmoregulatory and enzymatic adjustments that sustain its osmotic homeostasis in response to salinity acclimation, particularly useful in a challenging, variable salinity environment like the mangrove forest habitat.

## Acknowledgements

The authors thank the Instituto Chico Mendes de Conservação da Biodiversidade, Ministério do Meio Ambiente for authorization to collect *Ucides cordatus* under ICMBio/MMA permit #29594-9 to JCM. We also thank the Instituto Nacional de Ciência e Tecnologia para Adaptações da Biota Aquática da Amazônia (INCT-ADAPTA-II) with which this laboratory (FAL) is integrated, and the Rede de Camarão da Amazônia.

## Compliance with Ethical Standards

This study complies with all institutional, Brazilian and international guidelines on the use of invertebrate animals in scientific research.

## Author contributions

Preparation of biological material, and data collection and analyses were performed by Cintya M. Moraes, Leonardo M. Fabri, Malson N. Lucena, Rogério O. Faleiros, John C. McNamara and Carlos F.L. Fontes. The first draft and subsequent versions of the manuscript were written by Francisco A. Leone, John C. McNamara, Daniela P. Garçon and Leonardo M. Fabri. All authors participated in subsequent versions and read and approved the final version of the manuscript.

